# High-Fidelity Simulation of Cell Adhesion and Flow with Coarse Grain Modelling: A Numerical and Experimental Approach

**DOI:** 10.1101/2025.09.22.677682

**Authors:** Rajat Mishra, Saumya Jaiswal, Amit Arora, Abhijit Majumder

## Abstract

A coupled CFD-DEM framework was developed to investigate red blood cell (RBC) transport and cellular adhesion in constricted microchannels, with a primary objective of reducing computational cost while preserving physiological accuracy. Initially, numerical softening factors were implemented, allowing greater overlap between particles to expedite computations, with an optimal value of 0.03 selected. Adhesion forces between RBCs and the channel wall were incorporated, with a minimum effective distance of 0.4□µm identified as computationally feasible. However, RBC–RBC adhesion led to artificial clustering and was therefore omitted. Coarse-graining methods were applied up to a factor of 2 to further reduce simulation time, with a value of 1.8 chosen based on consistent cell-free layer (CFL) thickness and drag force characteristics. To correct for the increased drag in coarse-grained models, rolling resistance (Type C) with a coefficient of 0.8 was employed, bringing deviations within 10%. The validated model was then extended to simulate the detachment of adherent C2C12 myoblasts under physiological flow. Experimental validation showed increasing cell retention with seeding time, and simulations reproduced this trend with <5% deviation. The resulting Detachment Force Ratio (DFR) at 6 hours indicates near-equivalence between adhesion and hydrodynamic forces, establishing the model’s robustness for cell-substrate interaction studies in microfluidics.

## 1. Introduction

The dynamics of cellular suspensions in microfluidic environments are important for understanding a variety of physiological and biomedical phenomena, including blood flow, immune cell trafficking, and drug delivery [1,2]. In particular, the behavior of cells such as red blood cells (RBCs) and adherent cell types like myoblasts within confined geometries has implications for disease diagnostics, tissue engineering, and lab-on-a-chip device design [3–6]. Modeling such flows, however, presents unique challenges due to the need to resolve complex intercellular and fluid-structure interactions under low Reynolds number conditions [7].

Computational Fluid Dynamics (CFD) coupled with the Discrete Element Method (DEM) has emerged as a robust framework for simulating multiphase cell suspensions in microchannels [8,9]. This approach enables the tracking of individual cells as discrete particles while resolving fluid flow fields through the Navier–Stokes equations[10,11]. Although DEM-based techniques can capture detailed interactions between cells and the surrounding fluid, simulations often become computationally intensive when anatomically accurate cell shapes, deformability, or large cell populations are considered[12]. Consequently, many studies approximate cells as rigid particles and limit inter-particle physics, thereby sacrificing biophysical fidelity for computational tractability. Arya et al. [13] developed a numerical model to analyze flow of magnetic particles during drug delivery for cardiovascular diseases in carotid bifurcating artery. Magnets have been placed at four different places for different diameter of the magnetic particles and analysis had been made through velocity and shear profiles in order to analyze interaction at the desired locations. Quanyu et al. [14] developed a numerical model for the blood flow in human artery. The work modeled arterial flow in the human arm using Ansys Fluent, focusing on velocity, pressure, and shear stress in major arteries such as the brachial, radial, and ulnar. The work highlights key hemodynamic features at bifurcations and supports the clinical relevance of the radial artery, especially for noninvasive pulse monitoring.

To mitigate the above mentioned limitations with numerical modeling, recent efforts have explored numerical softening to mimic deformability, coarse-grained modeling (CGM) to reduce particle count, and simplified adhesion models to capture bonding effects between cells and walls or between cells themselves[15–17]. While these strategies offer partial solutions, an integrated CFD-DEM framework that balances computational efficiency with physical accuracy, especially for cases involving both non-adherent and adherent cell types, remains underdeveloped.

In the present study, we develop and validate a generalized CFD-DEM framework that models cell suspensions as rigid, discrete particles while systematically incorporating computational and physical enhancements to improve realism. We begin by investigating the flow of non-adherent RBCs through a constricted microchannel, incorporating numerical softening to allow controlled overlap and simulate the mechanical effects of cell deformability. An adhesion model is then introduced to simulate interaction forces between RBCs and the channel wall, as well as between RBCs, with a parametric evaluation of adhesion distance and its influence on clustering and cell-free layer (CFL) formation.

Subsequently, a coarse-graining strategy based on Bierwisch-type scaling is implemented to reduce computational cost, and its effects are corrected using rolling resistance models to preserve drag force accuracy[18]. The framework is extended to anatomically representative biconcave RBCs and finally adapted to simulate the adhesion behavior of C2C12 myoblast cells seeded on a collagen-coated microfluidic device. In these simulations, cell adhesion strength is calibrated against experimental detachment ratios (DR), defined as the ratio of cells remaining after flow to those initially seeded. A force-based detachment force ratio (DFR), representing the balance between adhesive and hydrodynamic forces, is also introduced to quantify detachment phenomena.

This comprehensive framework offers a scalable and physically grounded approach to simulating complex cellular interactions in microfluidic flows, providing new insights into the hydrodynamics of both non-adherent and adherent cells. The methodology and findings are relevant to applications in vascular modeling, mechanobiology, and organ-on-chip platforms [19–21].

## 2. Methodology

### 2.1 CFD-DEM coupling strategy

A single way coupled CFD-DEM model has been developed to analyze the flow of cells inside the channel. The equations of mass and momentum have been solved for fluid phase by classical Navier Stokes equation and are given by [22,23] :-

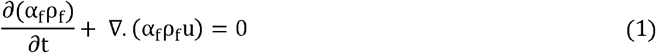

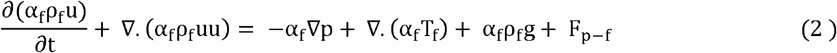

Where, α_f_ is the fluid volume fraction, ρ_f_ is the fluid density, p is the pressure, u is the fluid phase velocity vector, T_f_ is the stress tensor of the fluid phase.

Cells have been represented by solid discrete particles and have been tracked by solving Eulers first and second laws of motion that govern translational and rotational motion of cells. The translational motion is shown by Eq. (3) and rotational motion is shown by Eq. (4).

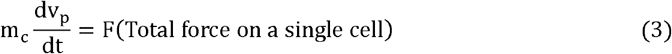

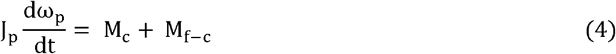

Where J_p_ is the moment of inertia, M_c_ is the net torque generated by the tangential forces that leads to rotation of cell, M_f-c_ is the additional torque that is generated by fluid phase velocity gradient.

Now in the translation motion equation, the total force on a single cell account for contact forces (F_c_) with other cells and contact force when it comes in contact with boundary of channel. The weight of the cell which is mg, where m is mass of a single cell and g is acceleration due to gravity. Also because of the fluid, cells feel the forces which is F_f-c_ and is forces on a cell because of the fluid. This accounts for drag force (F_D_), pressure gradient force (F_Δp_) and virtual mass force (F_VM_). The force balance on a single cell is represented in pictorial form in Fig. 1. Considering total force on a cell the mathematical description for the translation motion is shown below: -

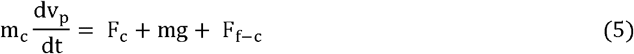

**Fig. 1.**
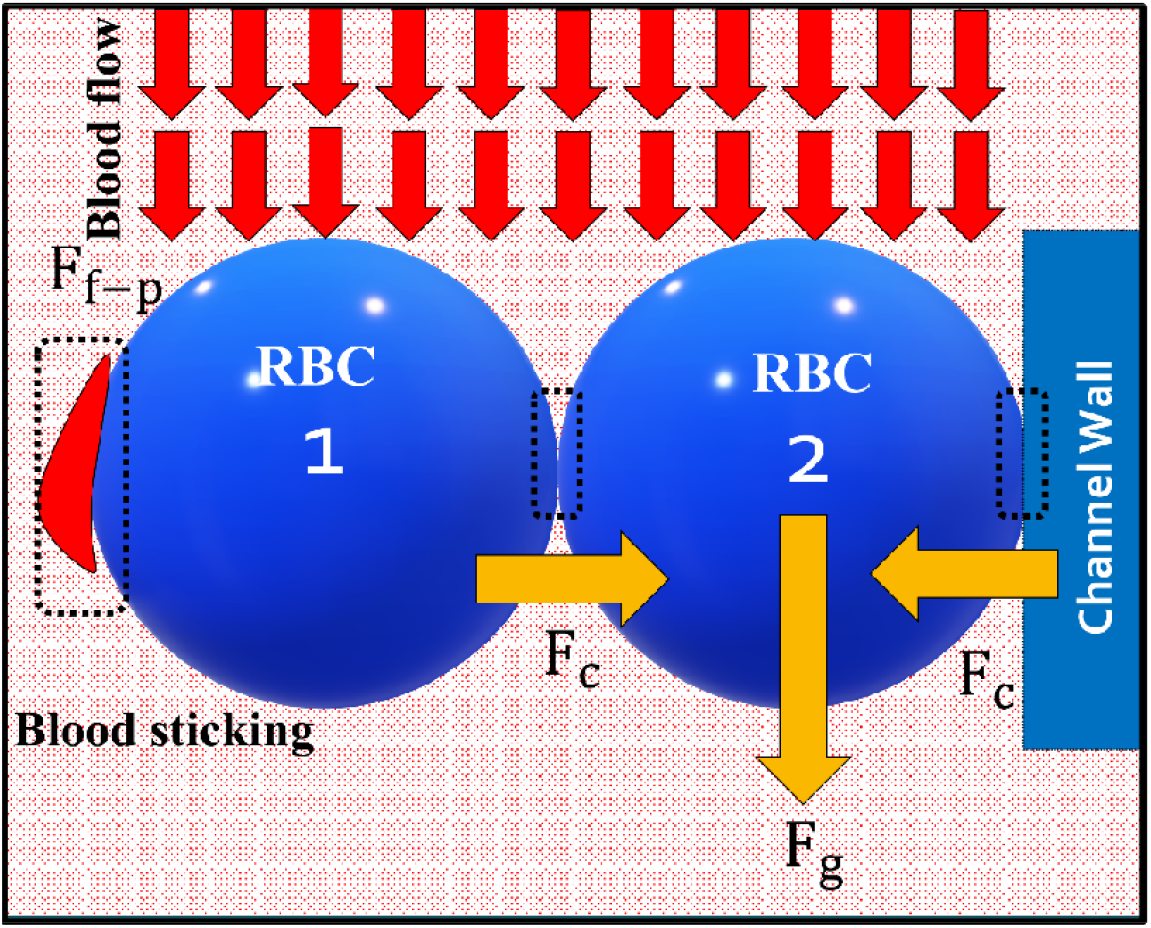
Image showing force balance on a flowing RBC cell inside constricted microfluidic device. The shown section is a very small section inside the channel considering force by fluid, channel wall and other RBCs on it.

The contact force is accounted by contact force models which evaluates tangential and normal contact forces. The hertzian spring dashpot model is used to evaluate normal contact force (F_n_) and coulomb limit model is used to evaluate tangential contact force (F_τ_) [24,25]. The mathematical description for the hertzian spring dashpot model is given in Eq. (6).

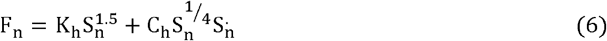

Where, K_h_ is the stiffness coefficient and C_h_ is the damping coefficient and mathematically represented in Eqs. (7), (8), respectively. S_n_ is contact normal overlap and S_n_ is time derivative of contact normal overlap.

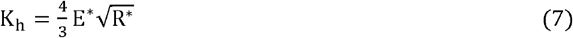

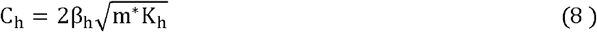

E^∗^ is the effective young modulus and R^∗^is the effective radius and is defined as below: -

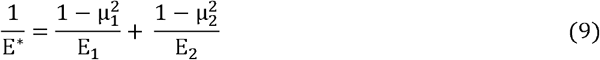

µ_1_, µ_2_ are the poisons ratio and E_1_, E_2_ are the young modulus for the contacting particles.

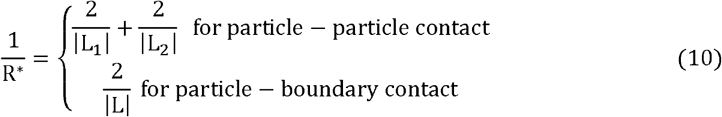

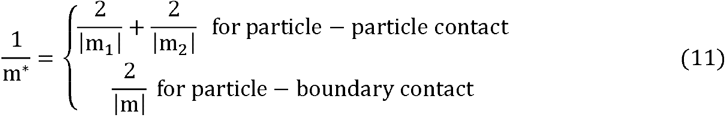

L_1_ and L_2_ are the size of the contacting particles and m_1_, m_2_ are the mass of the contacting particles.

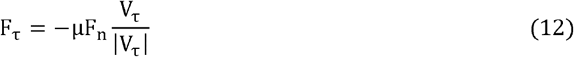

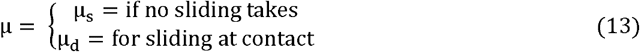

Now the term F_f-c_ is a summation of different forces which is mathematically represented below: -

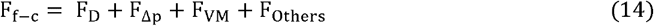

The forces include drag forces which have been evaluated by considering Huilin and Gidaspow drag force model for which the mathematical expression is shown in Eq. (15) [26]. This drag law transitions between Ergun and Wen drag laws by giving weightage to both drag laws through blending parameter φ [27,28]

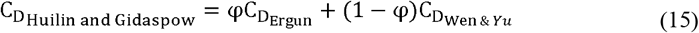

This parameter φ is a function of fluid volume fraction, which is defined as: -

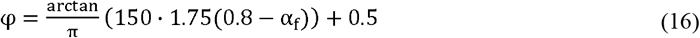

Ergun experiments were on fluid flow through a dense column of granular material and is valid for higher particle concentration (α_s_ ≥ 0.20). Pressure losses happened because of kinetic and viscous energy losses, for which the derived expression is shown below.

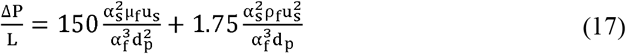

Where L is the height of the bed, Δp is pressure loss, d_p_ is the effective particle diameter and u_s_ is the superficial velocity.

Wen and Yu model is valid for low particle concentrations (α_s_ < 0.20). Expression for drag coefficient is given in equation based on experimental analysis by Gidaspow on fluidized beds.

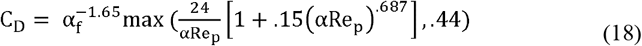

Where α_f_ is fluid volume fraction and Re_p_ is relative Reynold’s number for particles defined as: -

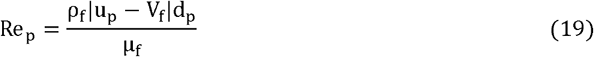

ρ_f_ is the fluid density, µ_f_ is fluid viscosity, d_p_ is particle diameter, *u*_*p*_ is particle velocity and V_f_ is fluid velocity.

Virtual mass force is evaluated by Ishi and Mishima’s model [29] for which the mathematical description is shown below:

is the virtual mass coefficient which is defined as below: -

### 2.2 Geometry and Flow conditions for RBC

A constricted microchannel with a hyperbolic contraction profile was selected to investigate blood flow dynamics and cell-free layer (CFL) formation, as illustrated in Fig. 2(a). The channel geometry was constructed using ANSYS SpaceClaim 2021 R1, based on dimensions reported in the experimental study conducted by [30]. A mesh-independence study was performed to ensure numerical accuracy, and the final meshed domain comprised 1,381,884 elements and 1,463,440 nodes. The mesh was generated using tetrahedral elements with a minimum element size of 0.05 mm, growth rate of 1.2, and a transition ratio of 0.272, as shown in Fig. 2. A steady inlet velocity of 2.5 mm/s was imposed at the channel entrance, replicating experimental flow rates. Red blood cells were continuously injected at the inlet and tracked throughout the channel. RBCs were modeled as rigid, non-deformable particles with an equivalent spherical diameter, enabling simplification of particle dynamics while maintaining the accuracy of CFL estimation.

**Fig. 2.**
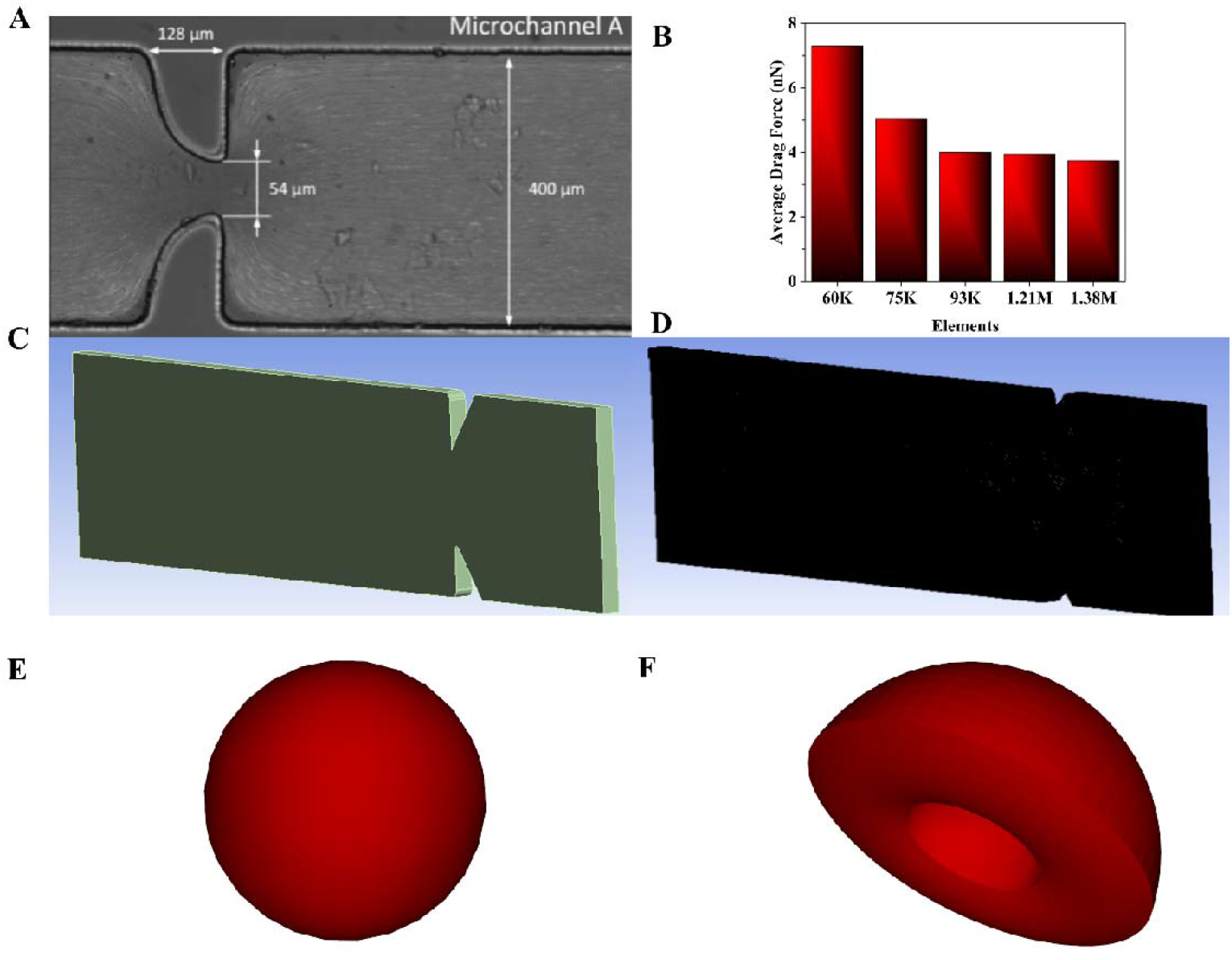
Channel geometry and particle details. **(A)** Dimensions of the constricted microchannel considered as a reference to analyze flow of RBCs for the developed coupled CFD-DEM model [30]. **(B)** Mesh independent analysis by comparing average drag force with number of elements. **(C)** CAD geometry of the microchannel used in the simulation. **(D)** Meshing of the microchannel. **(E)** Spherical shaped RBC. **(F)** Bi-concave shaped RBC

Given the low Reynolds number (Re < 1), the flow was modeled as laminar. A coupled CFD–DEM (Computational Fluid Dynamics-Discrete Element Method) framework was implemented to simulate cell-fluid interactions. A one-way coupling approach was employed, assuming that the fluid affects cell motion, but the reverse interaction is negligible due to the low RBC concentration (hematocrit 5%).

In this two-phase model, Dextran 40 (Dx40) was used to simulate the plasma phase, while RBCs constituted the discrete solid phase. The fluid and particle properties, along with DEM calibration parameters, are provided in Table 1.

**Table 1:** Properties of the RBC and calibrated values of the DEM parameters used in the simulation.

|  |  |  |
| --- | --- | --- |
| Dextran 40 Density | 1042 kg/m<sup>3</sup> | [30] |
| Dextran 40 Viscosity | 0.0052 kg/ms | [30] |
| RBC Density | 1185 kg/m<sup>3</sup> | [30] |
| RBC Young Modulus | 7 KPa | Calibration |
| RBC Diameter | 8 $\mu\text{m}$ (Equivalent sphere) | [30] |
| RBC Poisson's Ratio | 0.40 | Calibration |
| Static Friction | 0.60(C-W), 0.60(C-C) | Calibration |
| Dynamic Friction | 0.57(C-W), 0.55 (C-C) | Calibration |
| Tangential Stiffness Ratio | 1 | Calibration |
| Restitution Coefficient | 0.30 | Calibration |
| Drag Law | Huilin & Gidaspow | Calibration |

### 2.3 Numerical Softening Factor

In the present study, cellular dynamics in microchannel flow, comprising both non-adhering cells (e.g., red blood cells, RBCs) and adhering cells (e.g., C2C12 myoblasts) are modeled using the Discrete Element Method (DEM), where cells are represented as rigid particles. Although this rigid-body assumption simplifies computation, real biological cells are compliant and capable of deformation. To approximate this deformability within a rigid DEM framework, a numerical softening factor is introduced. This factor modifies the particle-particle stiffness in the contact model, allowing controlled overlap during cell-cell or cell-wall interactions, thereby emulating mechanical compliance. In the present model Eq. (22) shows the mathematical representation for calculation of the time step.

Lowering the stiffness not only mimics the deformation behavior of soft cells during contact but also increases the critical DEM time step, which scales inversely with the square root of the stiffness. This results in a substantial reduction in computational cost, particularly advantageous when simulating large numbers of non-spherical particles or incorporating adhesion mechanics. A hysteretic linear spring contact model is used to capture the loading-unloading behavior of interacting cells, and the critical time step is derived based on the reduced stiffness, cell mass, and damping characteristics to ensure numerical stability. This modeling strategy provides a computationally efficient yet physically representative framework for simulating the hydrodynamic and adhesive behavior of both suspended and surface-adherent cells under flow conditions in microfluidic systems.

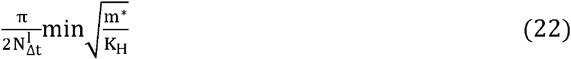

### 2.4 Adhesion Force Modeling

The interaction of biological cells such as red blood cells (RBCs) or adherent myoblasts (e.g., C2C12) with their surrounding environment, including neighbouring cells and microchannel walls, generates adhesion forces that significantly influence cellular behavior under flow [31]. These adhesive interactions are crucial in governing aggregation, clustering, and retention phenomena. In non-adherent systems like RBC suspensions, adhesion plays a role in margination and the formation of transient aggregates (e.g., rouleaux), whereas in adherent systems, it directly determines the extent of cell detachment or stability under shear flow[32].

To capture these effects, an adhesion force model is incorporated within the DEM framework. The model, defined mathematically in Eq. (23), activates when the surface-to-surface distance between two particles (or a particle and a wall) falls below a defined cutoff. The force is typically a function of this separation distance and includes tuneable parameters such as adhesion strength and interaction range.

Introducing adhesion is especially critical when numerical softening and coarse-grained modeling (CGM) are employed, as these techniques can reduce the natural contact duration or dilute intercellular interactions. Adhesion modeling ensures that essential behaviors such as clustering of RBCs or detachment of C2C12 cells are preserved despite numerical simplifications. The influence of adhesion is systematically analyzed in combination with softening and CGM to maintain consistency with experimental detachment ratios and known biological behaviors.

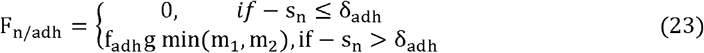

Where, F_n/adh_ is contact normal overlap, s_n_ is contact normal overlap, m_1_ and m_2_ are mass of particles in contact, g is the acceleration due to gravity, f_adh_ is force fraction, and δ_adh_ is adhesive distance.

### 2.5 Coarse Grain Modeling

To further alleviate the computational burden of simulating large populations of biological cells under flow, a Coarse-Grained Modeling (CGM) strategy is employed within the CFD-DEM framework. In this approach, a group of real cells is represented by a single, larger virtual particle, thereby reducing the total number of discrete elements in the simulation domain. This strategy is particularly useful for capturing large-scale behaviors of non-spherical or adherent cells—such as red blood cells (RBCs) or C2C12 myoblasts while maintaining key mechanical and hydrodynamic interactions [15,33]. The coarse-graining is implemented by scaling up the particle size using a coarse-graining factor (Λ), which necessitates appropriate adjustments to contact forces, drag coefficients, and other interaction parameters to retain physical consistency. Fig. 3. shows the coarse grain scaling for scaling factor of 2 for RBC cells. The set of eight collisions is replaced by a single set of collision and mass is same for comparison of both the systems.

**Fig. 3.**
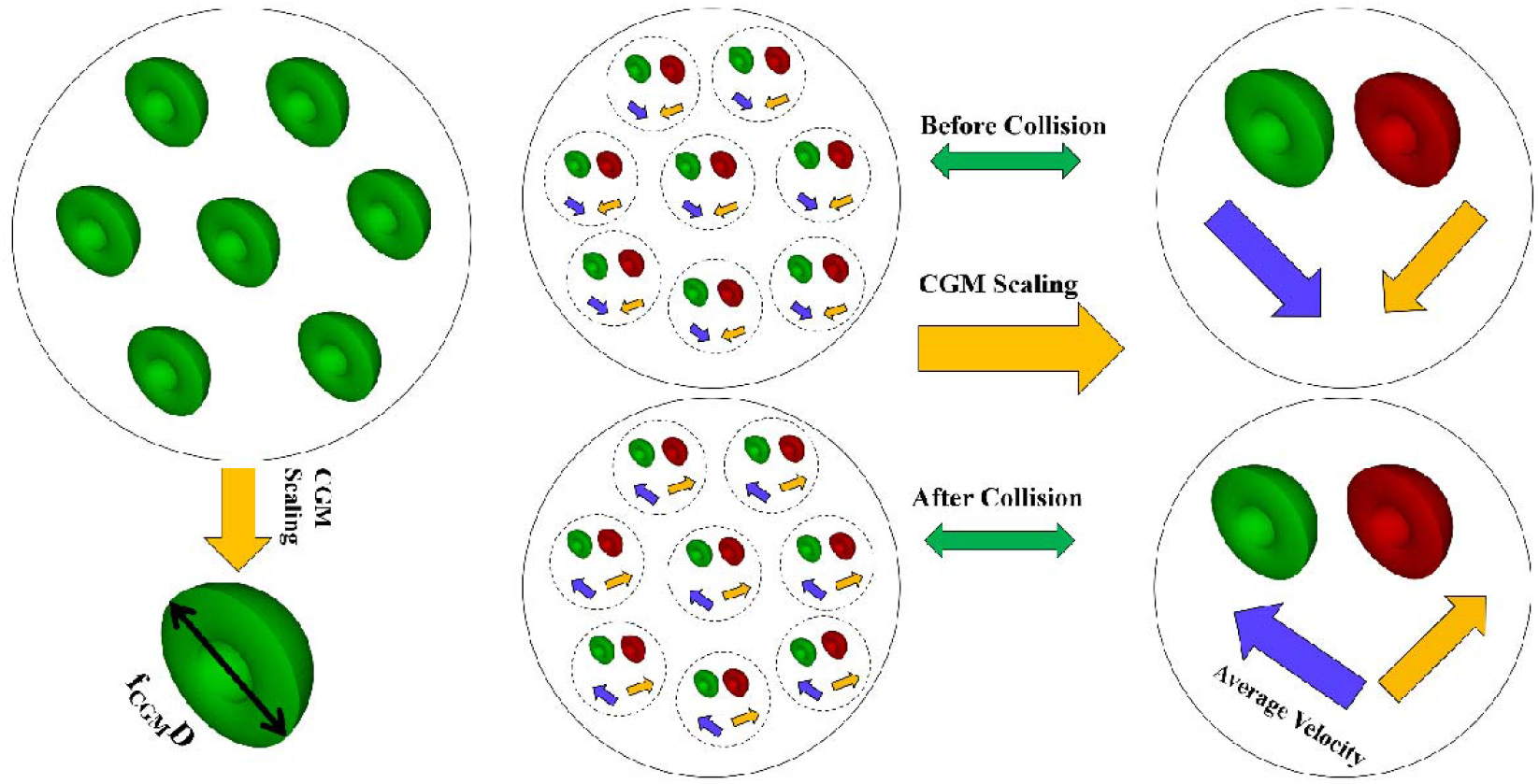
Principle of the Coarse Grain Model with scaling factor f_CGM_. The example shows scaling with f_CGM_ as 2, that lead to set of eight collisions between RBCs (coloured in green and red) being replaced by set of one collision. The mass is conserved before and after scaling so the kinetic energy.

Multiple values of the coarse-graining factor are systematically tested to evaluate their influence on critical flow features. Specifically, average drag force and cell-free layer (CFL) thickness are computed for each case and benchmarked against the baseline (unscaled) simulation. These metrics serve as proxies for momentum exchange and near-wall depletion behavior, respectively. For adherent cells like C2C12, additional care is taken to ensure that coarse-graining does not suppress essential adhesion dynamics or alter detachment trends under shear.

While coarse-graining offers substantial reductions in computational time, it can introduce artifacts, such as artificial spikes in drag force or dampened collision dynamics. To address this, rolling resistance models are implemented, and their parameters are tuned to counteract exaggerated rotational and contact behaviors arising from enlarged particles. The combined effect of CGM, adhesion modeling, and numerical softening is analyzed to ensure that both physical realism and numerical stability are retained across biologically relevant flow scenarios.

The results, including relative errors in average drag force and CFL thickness across different values of Λ, are summarized in Table 2. These findings demonstrate that, with appropriate corrections, CGM can be a powerful tool for extending simulations to biologically meaningful time scales and domains without compromising essential physics.

**Table 2.** Details of the DR and FDR values in experiments and simulation for different seeding time of C2C12 cells on collagen matrix.

| Seeding Time (hr) | DR |  | FDR |
| --- | --- | --- | --- |
|  | Experiment | Simulation |  |
| 1 | 58.38% | 60.40 | 0.44/0.35/5.28pN |
| 2 | 73.60% | 75.60% | 0.66/.52/7.85 |
| 4 | 83.90% | 86% | 0.78/.62/9.36 |
| 6 | 92.40% | 94.80 | 0.96/.76/11.48pN |

### 2.6 Rolling resistance model

In order to correct the coarse grained model by reducing the average drag force values, rolling resistance models of constant moment Type A and Type C have been used. The mathematical description of the Type A model is shown below in equations [34].

Where,

µ_r_ is rolling resistance coefficient, F_n_ is contact normal force, F_n_ is particle angular velocity vector and r is particle rolling radius, ω is particle angular velocity vector.

Type C is viscous damping model having viscous term to account for the oscillations. The mathematical description of the rolling resistance models Type C is given below: -

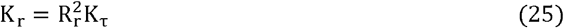

Where,

K_τ_ is tangential stiffness defined in Eq (7). R_r_ is the rolling radius given as below: -

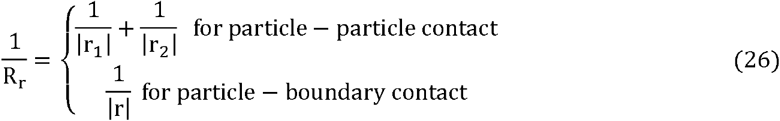

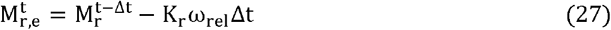

Where, 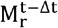 is the rolling resistance moment, Δt is the simulation timestep, 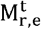 is the rolling resistance moment at the previous timestep.

### 2.7 Experimental Details

#### Device setup

Microfluidic devices were fabricated using polydimethylsiloxane (PDMS) mixed with a curing agent (Sylgard 184 Silicone Elastomer, Dow Corning) in a 10:1 weight ratio. The mixture was thoroughly degassed in a desiccator to eliminate air bubbles and then poured onto a master mold. The mold was subsequently placed in a hot-air oven at 70□°C for 2 hours to cure the PDMS. After curing, the PDMS devices were carefully peeled off from the mold, and 1 mm biopsy punches were used to create inlet and outlet ports as illustrated in Fig. 4. For flow experiments, a syringe pump (Company name and model to be added) was connected to the inlet via 1 mm inner-diameter tubing. The outlet tubing was directed into a collection reservoir. The entire flow setup with the tubing and microfluidic devices was placed inside a CO□ incubator maintained at 37□°C, while the pump was positioned externally above the incubator, as shown in Fig. 4(b), (c) and (d).

**Fig. 4.**
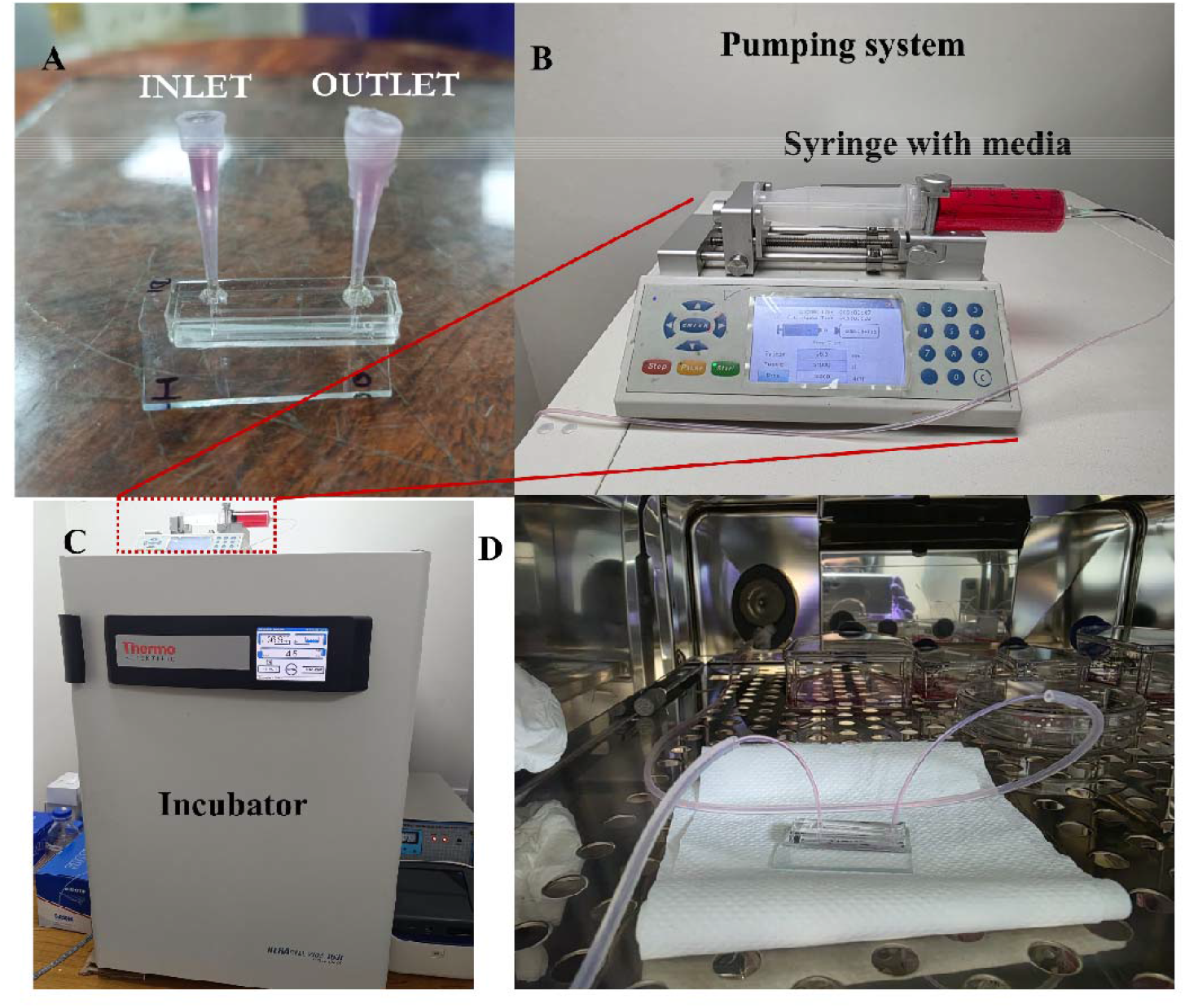
Images showing experimental setup and device used for the flow analysis of adhered cells. **(A)** Microfluidic device having channel with dimension of 1*1*30 mm having inlet and outlet for media. **(B)** Pump syringe setup filled with media. **(C)** Incubator with syringe pump system placed above it. **(D)** Microfluidic device placed inside the incubator with pipes attached at inlet and outlet for flow of the media.

#### Cell culture

Mouse myoblast cell line, C2C12 was seeded in the devices to study cell adhesion over time. Prior to seeding, the inner surface of the microchannels was coated with collagen solution (25□μg/mL) and incubated to facilitate surface functionalization. A total of 15,000 cells per straight channel were seeded, and the devices were prepared to evaluate adhesion at 1, 2, 4, and 6 hours, respectively. Cells were cultured using DMEM High Glucose medium supplemented with 20% FBS. For flow-based deadhesion analysis, a 60 mL BD syringe with culture medium was used to maintain flow at a rate of 5□μL/s for 30 minutes. Pre and post-flow imaging and analysis were performed to quantify adherent versus detached cells which is discussed in detail in results and discussion section.

## 3. Results and Discussion

### 3.1 Calculation of cell free layer and average drag force

#### 3.1.1 For numerical softening factor case

To evaluate the impact of numerical softening on simulation accuracy and computational efficiency, various softening factors were applied: 1, 0.5, 0.1, 0.05, and 0.03. For each case, the transient and time-averaged drag forces acting on the red blood cells (RBCs) were computed over a simulation duration of 10 seconds. Fig. 5(a) presents the transient variation of the average drag force for each softening value, while Fig. 5(b) illustrates the corresponding average drag force over the full simulation period. The results indicate that the drag force remains largely unaffected by the reduction in stiffness. This is attributed to the fact that drag force calculations primarily depend on the projected surface area of the particles, which remains unchanged under softening since only the contact stiffness is altered—not the particle geometry or hydrodynamic exposure. Additionally, Fig. 5(d) displays simulation snapshots capturing the cell-free layer (CFL) development across the tested softening factors. The visual analysis confirms that CFL formation remains consistent, with negligible variation among the cases.

**Fig. 5.**
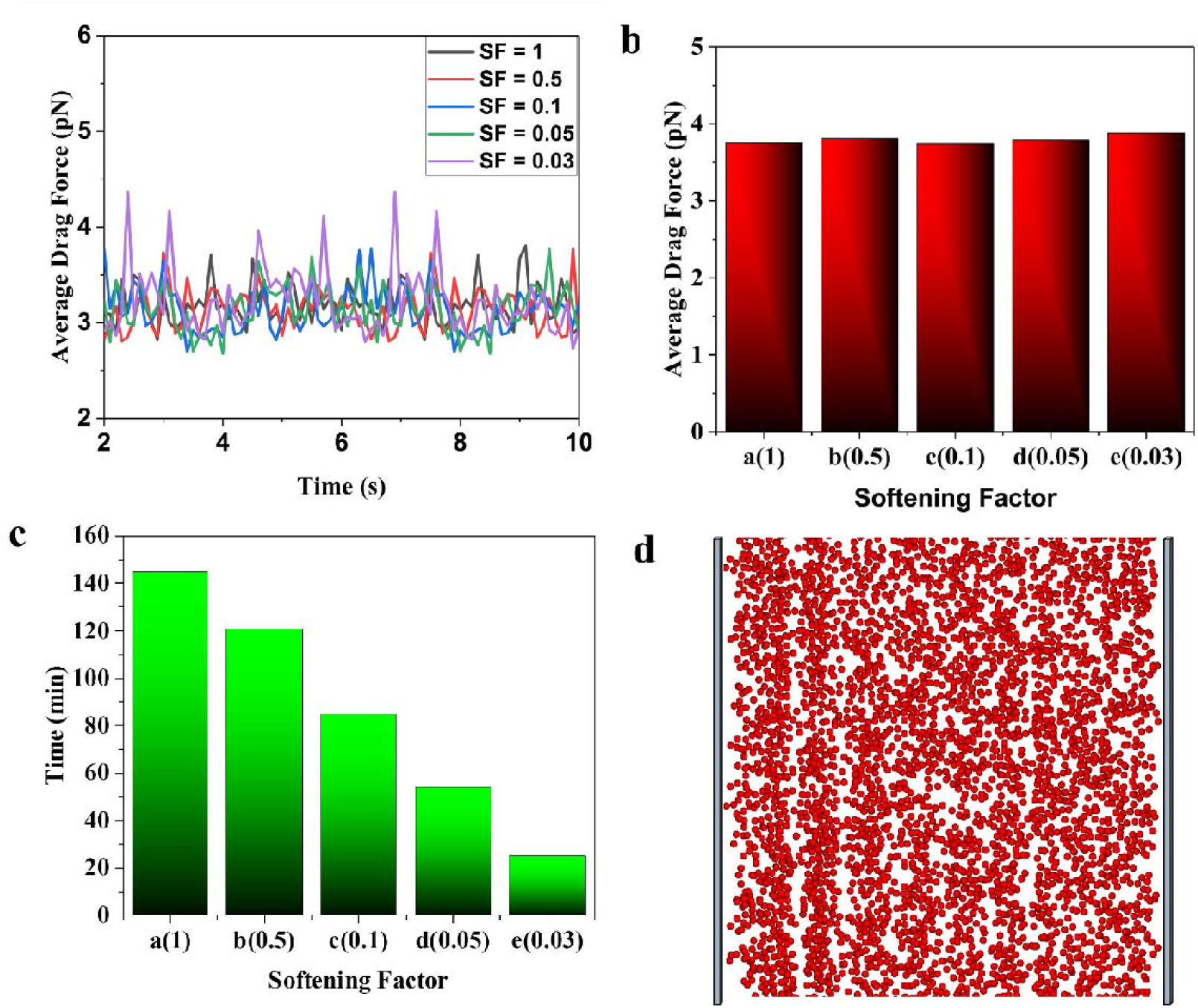
Postprocessing of the results for numerical softening factor case. **(a)** Transient plot for average drag force for different values of numerical softening factor. **(b)** Plot for average drag force values. **(c)** Computation time for different values of the softening factor considered in the simulation. **(d)** Snapshot of the simulation showing cell free layer formation

The computational performance, summarized in Fig. 5(c), demonstrates a substantial reduction in simulation time as the softening factor decreases, highlighting the benefit of using softened contact parameters, particularly for computationally intensive simulations involving nonspherical RBC geometries. However, further[35] reduction beyond SF=0.03 was restricted by the solver, as it led to excessive overlap between particles (exceeding ~17%), violating physical plausibility and numerical stability.

Based on these observations, a softening factor of 0.03 was selected for subsequent simulations. This value strikes a balance between maintaining accurate drag and CFL predictions while significantly reducing computational time.

#### 3.1.2 With adhesion force

The adhesion force model was incorporated by defining a minimum adhesive interaction distance, beyond which the adhesive force becomes inactive. This threshold distance was varied to study interactions both between RBCs and between RBCs and the channel wall. For RBC–RBC interactions, adhesive distances of 0.1, 0.2, and 0.3 µm were investigated. For RBC–wall interactions, the adhesive distance was varied from 0.1 to 0.4 µm.

Initially, only the RBC–RBC adhesive interactions were activated. Fig. 6(a) illustrates the average drag force, which increases with increasing adhesive distance. This is attributed to the stronger intercellular attraction leading to more frequent and prolonged contact events. Fig. 6(b) shows that CFL thickness also increases with larger adhesive distances, likely due to enhanced clustering near the center of the flow and reduced cell dispersion toward the walls. Subsequently, the analysis was repeated by activating only the RBC–wall adhesion as shown in Fig. 7. As seen in Fig. 7(c), the average drag force remains relatively unchanged with increasing adhesive distances. Furthermore, although a modest change in CFL thickness (~3 µm) was observed for the maximum adhesive distance (0.4 µm), it came at the cost of increased computation time.

**Fig. 6.**
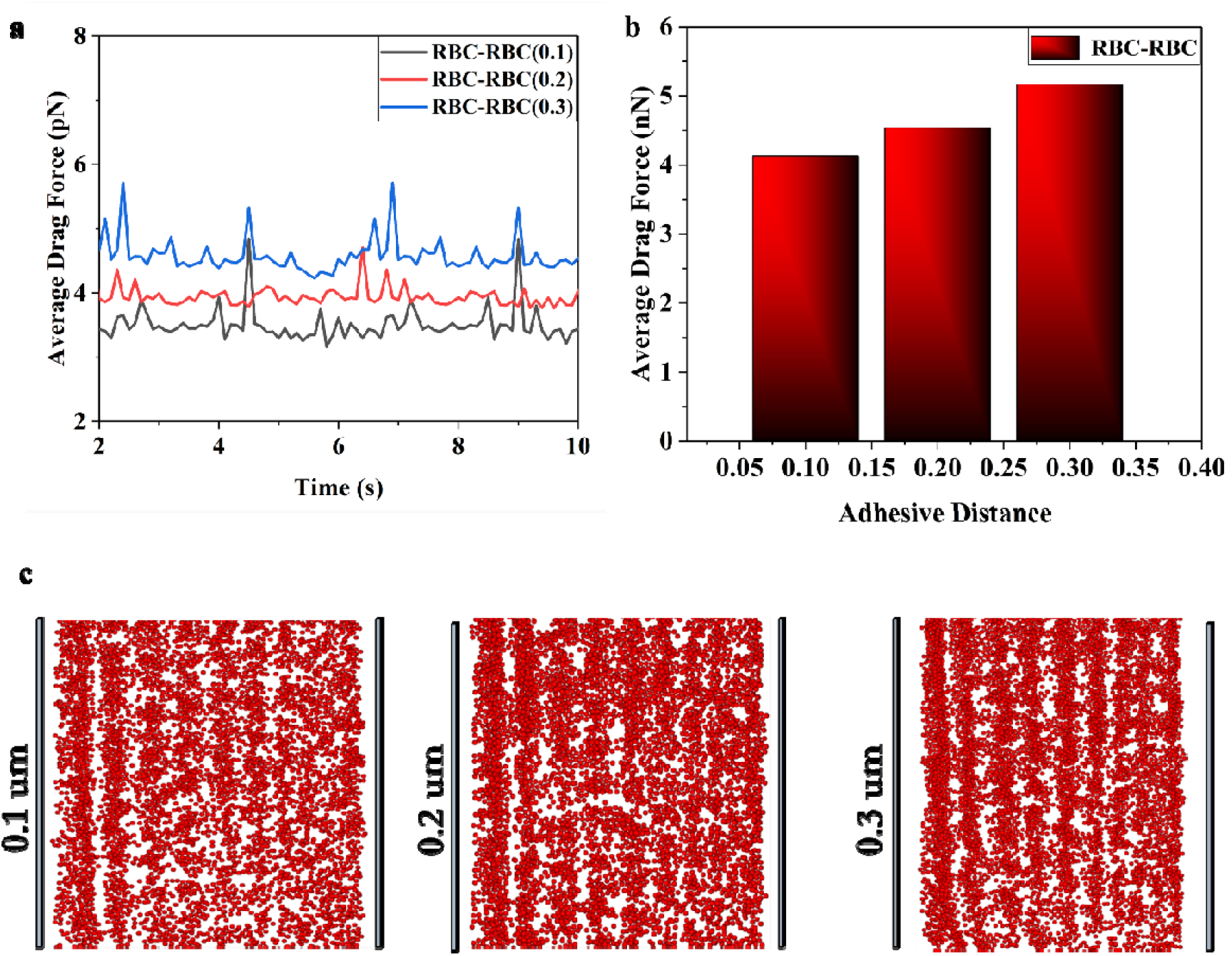
Postprocessing of the results for the case when adhesion force model is included between RBCs. **(a)** Plot for transient average drag force for different values of minimum adhesive force distance between RBCs. **(b)** Average drag force histogram plot **(c)** Snapshots of the simulation showing cell free layer formation for different values of minimum adhesive distance.

**Fig. 7.**
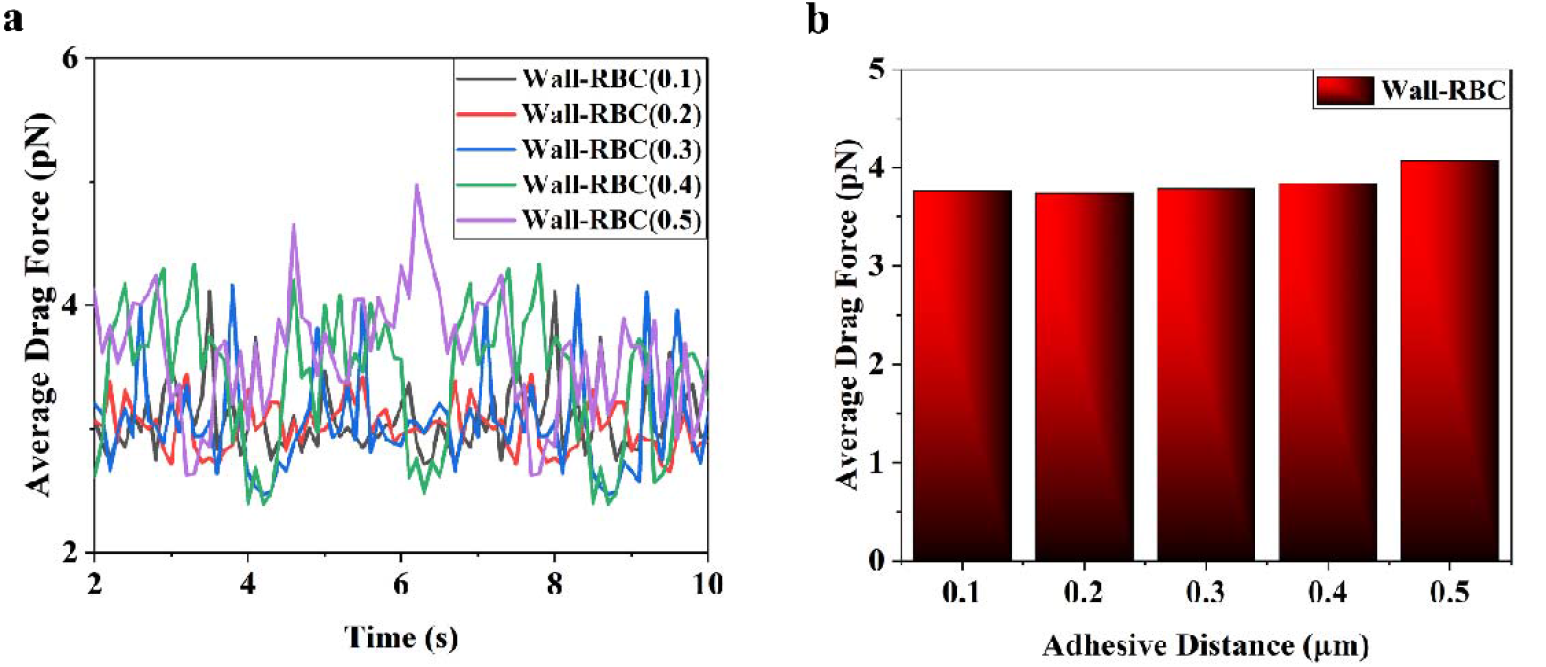
Postprocessing of the results for the case when adhesion force model is included between channel wall and RBCs. **(a)** Transient average drag force plot for different values of minimum adhesive force distance between RBCs and channel wall. **(b)** Histogram plot for the average drag force values

Finally, both RBC–RBC and RBC–wall adhesive interactions were considered simultaneously as shown in Fig. 8. The results confirm that the increase in drag force is primarily driven by RBC–RBC interactions, with RBC–wall adhesion having negligible impact on drag magnitude. Given that experimental studies report CFL thickness values in the range of ~5 µm, the observed increase in CFL due to RBC–RBC adhesion aligns with physical expectations.

**Fig. 8.**
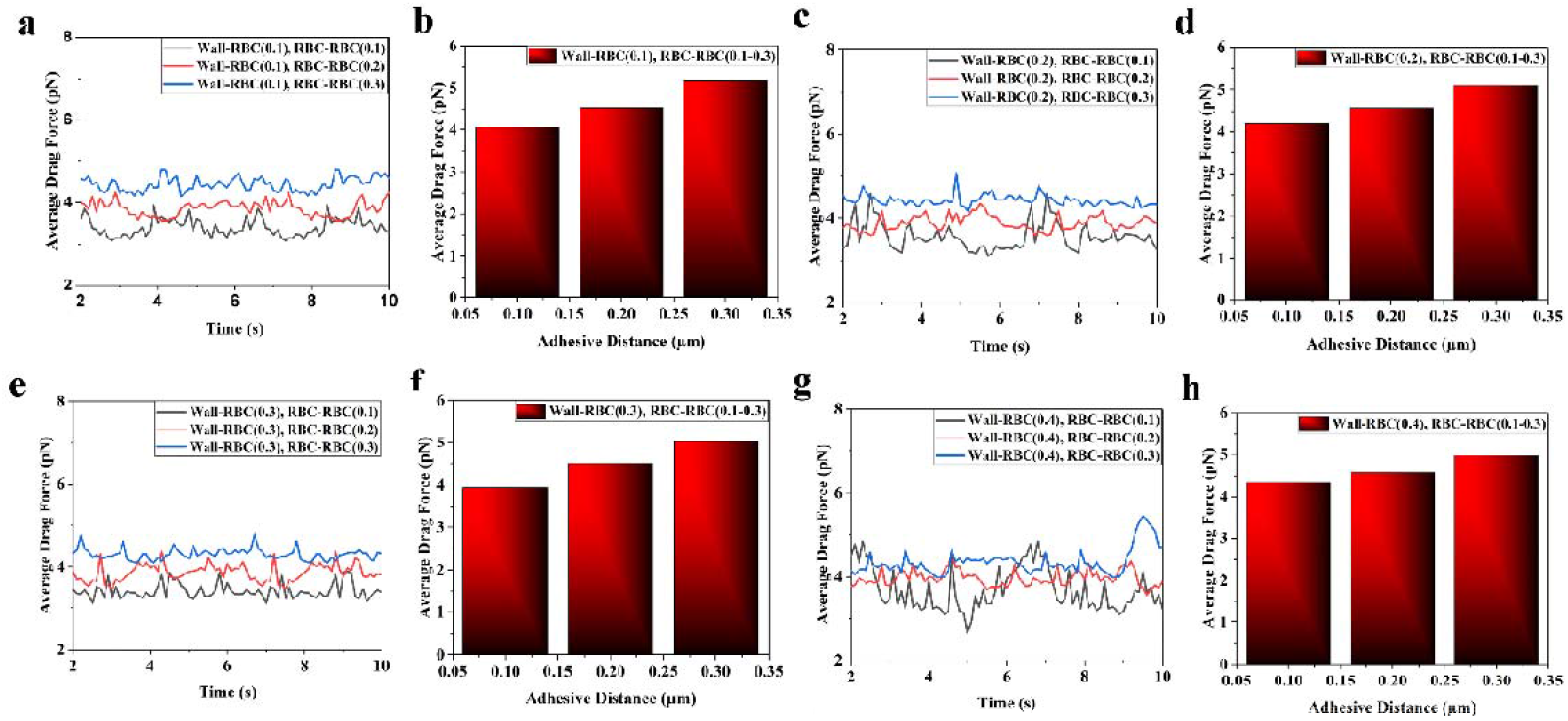
Postprocessing of the results for the case when adhesion force model is included between channel wall and RBCs. and between RBCs. Transient average drag force plot for different values of the minimum adhesive force distance between Wall-RBC and between RBCs. **(a)** 0.1 between wall and RBC **(b)** Histogram plot for the average drag force value. **(c)** 0.2 between wall and RBC **(d)** Histogram plot for the average drag force value. **(e)** 0.3 between wall and RBC **(f)** Histogram plot for the average drag force value. **(g)** 0.4 between wall and RBC. **(h)** Histogram plot for the average drag force value.

For subsequent analyses, only the RBC–wall adhesion with a minimum interaction distance of 0.4 µm was retained, as it can qualitatively represent RBC clustering and margination, although these effects are not the focus of the present study. This choice balances biological realism with computational efficiency while excluding the complexity introduced by strong intercellular adhesion

#### 3.1.3 With CGM

In order to evaluate the influence of coarse-graining on simulation performance and flow characteristics, CGM scaling factors ranging from 1.2 to 2.2 (in increments of 0.2) were considered. Fig. 9 presents the average drag force values for each CGM level. It is evident that as the CGM factor increases, the average drag force also increases, and the percentage deviation from the original unscaled case grows accordingly. This deviation is attributed to the reduction in particle number and increased particle size, which affects the force balance in the system. Fig. 9(b) illustrates the computation time corresponding to different CGM values. A clear reduction in simulation time is observed with increasing CGM factors, validating the benefit of CGM in improving computational efficiency. Fig 9(c) shows the histogram plot for average drag force value for different CGM parameters.

**Fig. 9.**
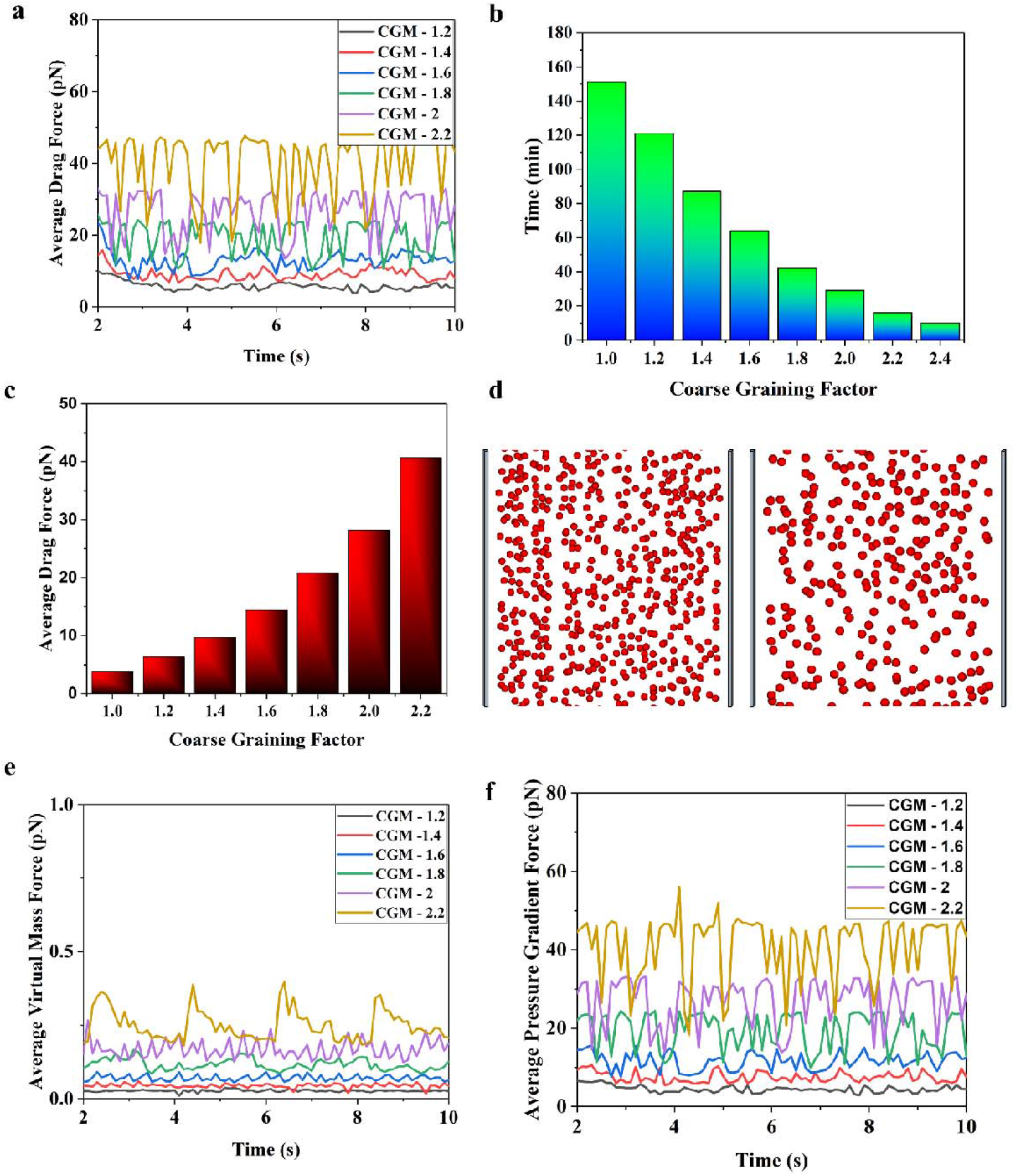
Postprocessing of the results for the coarse grain modelling case. **(a)** Transient plot for average drag force for different values of the coarse grain factor. **(b)** Histogram plot showing computation time for different values of coarse graining factor. **(c)** Histogram plot for average drag force value for different coarse grain value. **(d)** Snapshots of the simulation showing cell free layer formation for coarse graining factor of 1 and 1.8 and 2. **(e)** Transient plot for average virtual mass force for different values of coarse graining factor. **(f)** Transient plot for average pressure gradient force for different values of coarse graining factor.

Fig. 9(d) shows the cell-free layer (CFL) thickness as a function of CGM. Notably, no significant change in CFL is observed for CGM values up to 1.8, indicating that structural features of RBC distributions are well preserved. However, at CGM = 2.0, the number of RBCs becomes insufficient to reliably resolve the CFL, and drag force deviation becomes more pronounced. Thus, for subsequent simulations, a CGM value of 1.8 was selected as an optimal trade-off between accuracy and computational cost.

Additionally, Fig. 9(e) and 9(f) compares the average virtual mass force and pressure gradient force across different CGM values. While both forces exhibit an increasing trend with higher CGM, the virtual mass force remains substantially lower compared to drag and pressure gradient forces, indicating its relatively minor role in the overall force balance in this context.

### 3.2 Corrected drag force with rolling resistance models

The implementation of coarse-grained modeling (CGM) introduces deviations in key force metrics compared to the fully resolved case (CGM = 1.0). While a CGM factor of 1.8 was chosen to significantly reduce computation time, Fig. 10(a) shows that this scaling results in an approximate 350% increase in the average drag force, necessitating corrective measures to ensure physical accuracy.To address this discrepancy, rolling resistance models were incorporated into the CGM framework. Both Type A and Type C rolling resistance models were implemented to mitigate the overprediction of drag force observed in the coarse-grained system. Fig. 10(b) presents the average drag force values for varying rolling resistance coefficients using the Type C model. Although results for Type A are not shown for brevity, they were evaluated under similar conditions to ensure repeatability.

**Fig. 10.**
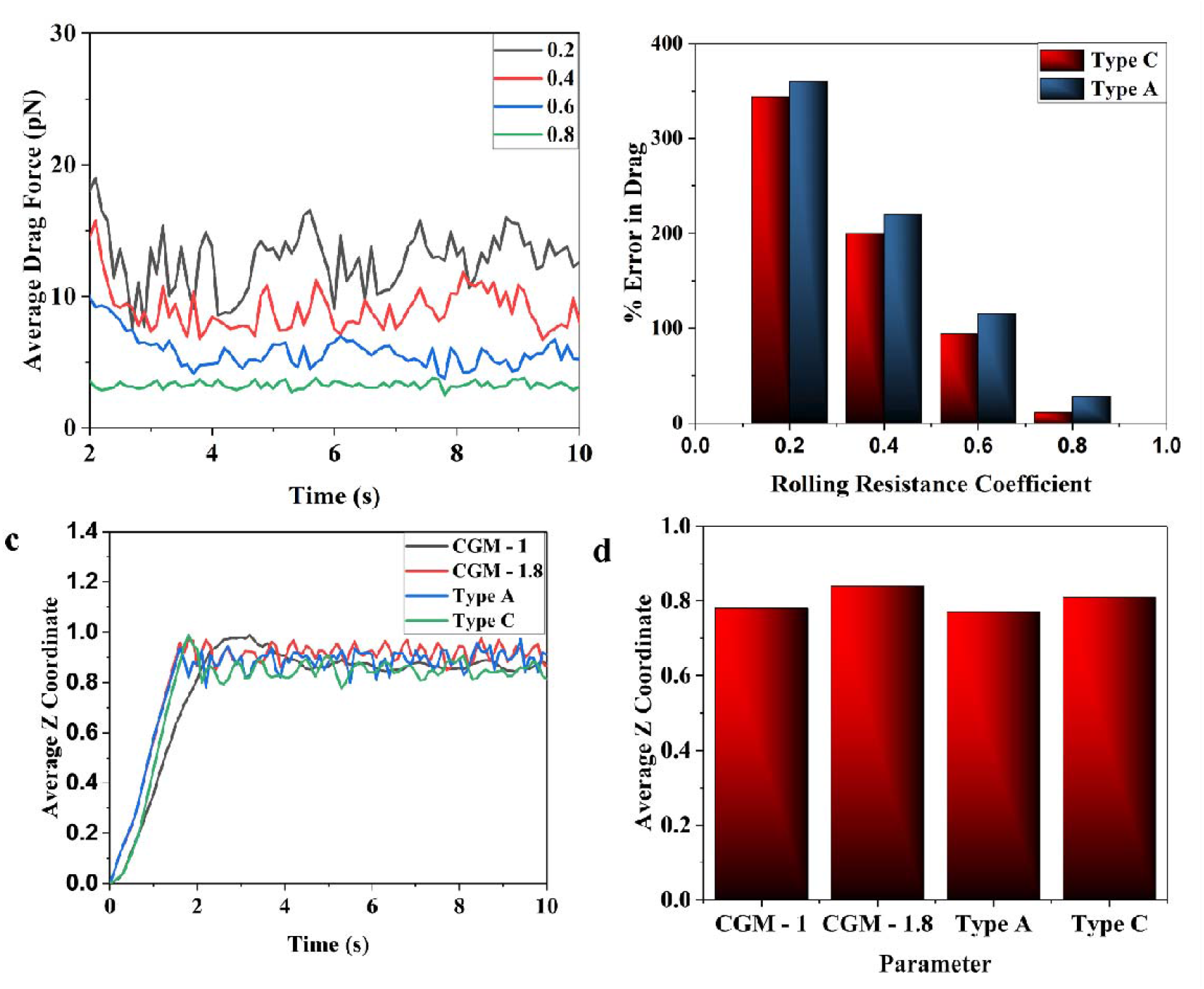
Postprocessing of the results for the corrected drag force when coarse grain model is incorporated by considering rolling resistance model. **(a)** Transient plot for the average drag force for different values of rolling resistance coefficients for rolling resistance model Type C. **(b)** Histogram plot showing comparison of average drag force between Type A and Type C rolling resistance model. **(c)** Transient plot for average Z co-ordinate for comparison between selected value of coarse graining factor and uncoarse-grained case and between Type C and Type A rolling resistance model at value of 0.8 rolling resistance coefficient. **(d)** Histogram plot for average Z coordinate for selected values of coarse graining factor and rolling resistance model.

The Type C model, which applies a constant resisting moment, proved more effective in reducing the average drag force than the Type A model, particularly for rolling resistance coefficients up to 0.8. Beyond this threshold, however, two adverse effects were observed: (i) a significant increase in computation time, and (ii) a reduction in the average Z-coordinate of RBCs, indicating altered flow behavior or vertical displacement as shown in Fig. 10(c) and (d).

Consequently, a rolling resistance coefficient value of 0.8 for the Type C model was selected as the upper limit for all further simulations, offering a balance between physical accuracy (in drag force correction) and computational efficiency.

### 3.3 Modeling for nonspherical RBC

Using the finalized values for the softening factor, adhesion force parameters, and coarse-grained modeling (CGM), simulations were conducted with biconcave (non-spherical) representations of RBCs. Fig. 11(a) illustrates a simulation snapshot highlighting the biconcave geometry. Fig. 11(b) presents a comparison of the average drag force between spherical and non-spherical RBC cases. The results show a slightly higher drag force for the biconcave RBCs, with average values of 24.2□pN for the non-spherical case and 21.8□pN for the spherical one. This increase can be attributed to the enhanced surface interaction and larger effective projected area presented by the biconcave shape. Fig. 11(d) shows the transient variation in average Z-coordinate, nondimensionalized by the channel length, for both cases. Due to higher drag, the non-spherical RBCs exhibit relatively greater displacement in the flow direction compared to their spherical counterparts.

**Fig. 11.**
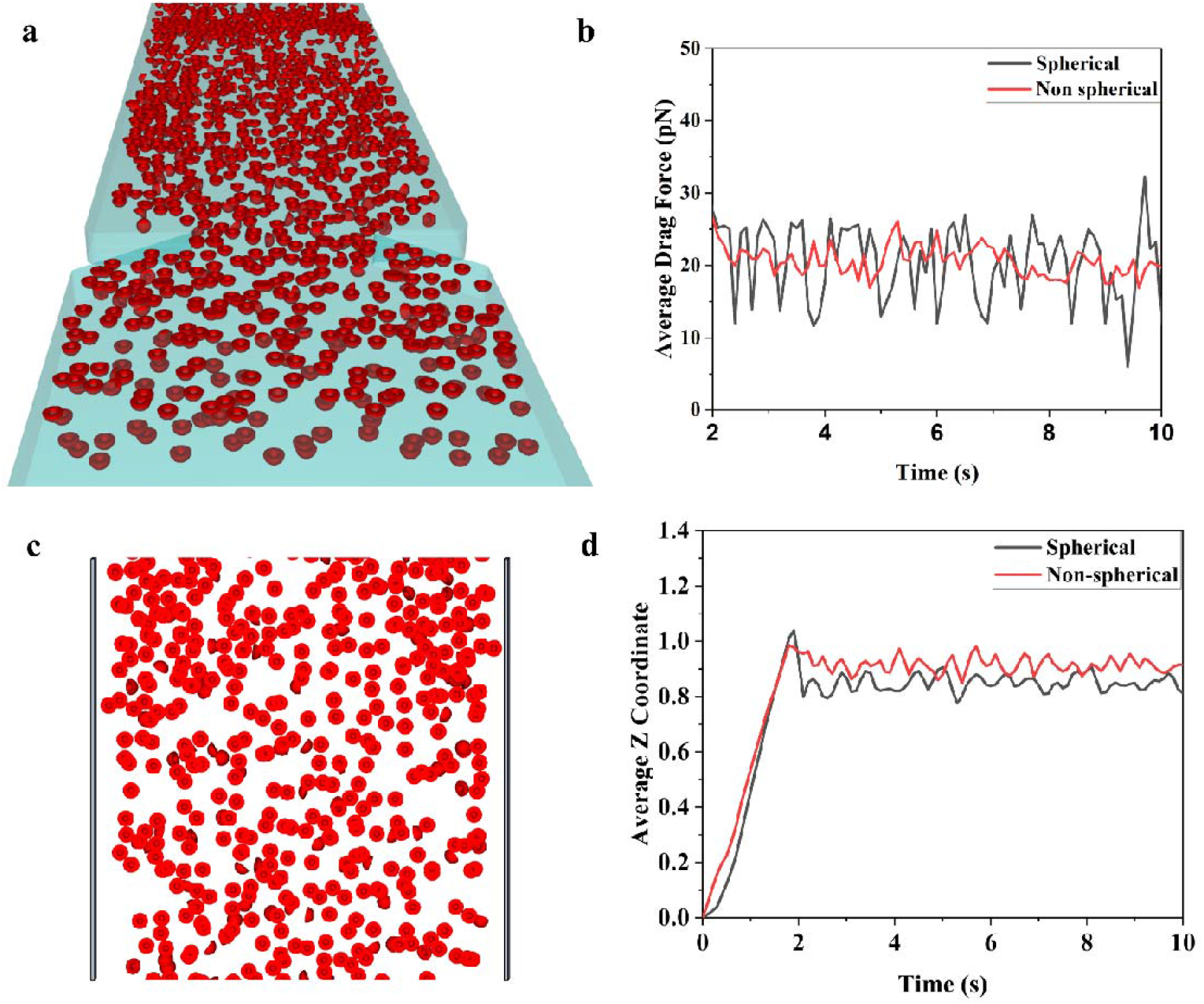
Postprocessing of the results when biconcave shape of RBC is considered. **(a)** Snapshot of the simulation showing flow of RBCs for bi-concave shape. **(b)** Transient average drag force plot showing comparison between spherical and non-spherical RBC. **(c)** Snapshot of the simulation showing cell free layer formation. **(d)** Transient plot for the average Z coordinate showing comparison between spherical and non-spherical RBC.

Despite the geometric complexity, no significant difference was observed in cell-free layer (CFL) thickness between the two cases, indicating that spherical approximation remains reasonable for CFL predictions. However, considering the fidelity of shape and interactions, the non-spherical model provides more accurate drag estimations.

Owing to the increased computational demand associated with complex contact and hydrodynamic interaction calculations for non-spherical particles, the 10-second simulation took approximately 6 days and 14 hours on the current computational setup.

### 3.4 Implementation for adhered cells

In the experimental setup, C2C12 cells were seeded onto collagen-coated microfluidic devices, and adhesion was assessed after incubation times of 1, 2, 4, and 6 hours. Flow analysis was subsequently performed using a syringe pump system connected to the microfluidic device. Microscopic images were acquired at specific regions of the channel, both before and after flow application, to evaluate the extent of cell detachment.

To replicate this scenario numerically, a microchannel with dimensions 1 × 1 × 30 mm was created, as shown in Fig. 12(a). The cells post-seeding was spherical particles and seeded onto the device using a two-step procedure. First, a volume fill region was defined within a segment of the channel, and particles were inserted under the influence of gravity and adhesion forces between the cells and the substrate. The seeding region was centered around a specified seed coordinate (Fig. 12(c)), around which the particles were initialized. A transient simulation of 0.5 seconds was conducted for cell deposition, resulting in the stable placement of 917 cells. The particle configuration at the final timestep of this simulation was extracted and used as the initial condition for subsequent flow simulations.

**Fig. 12.**
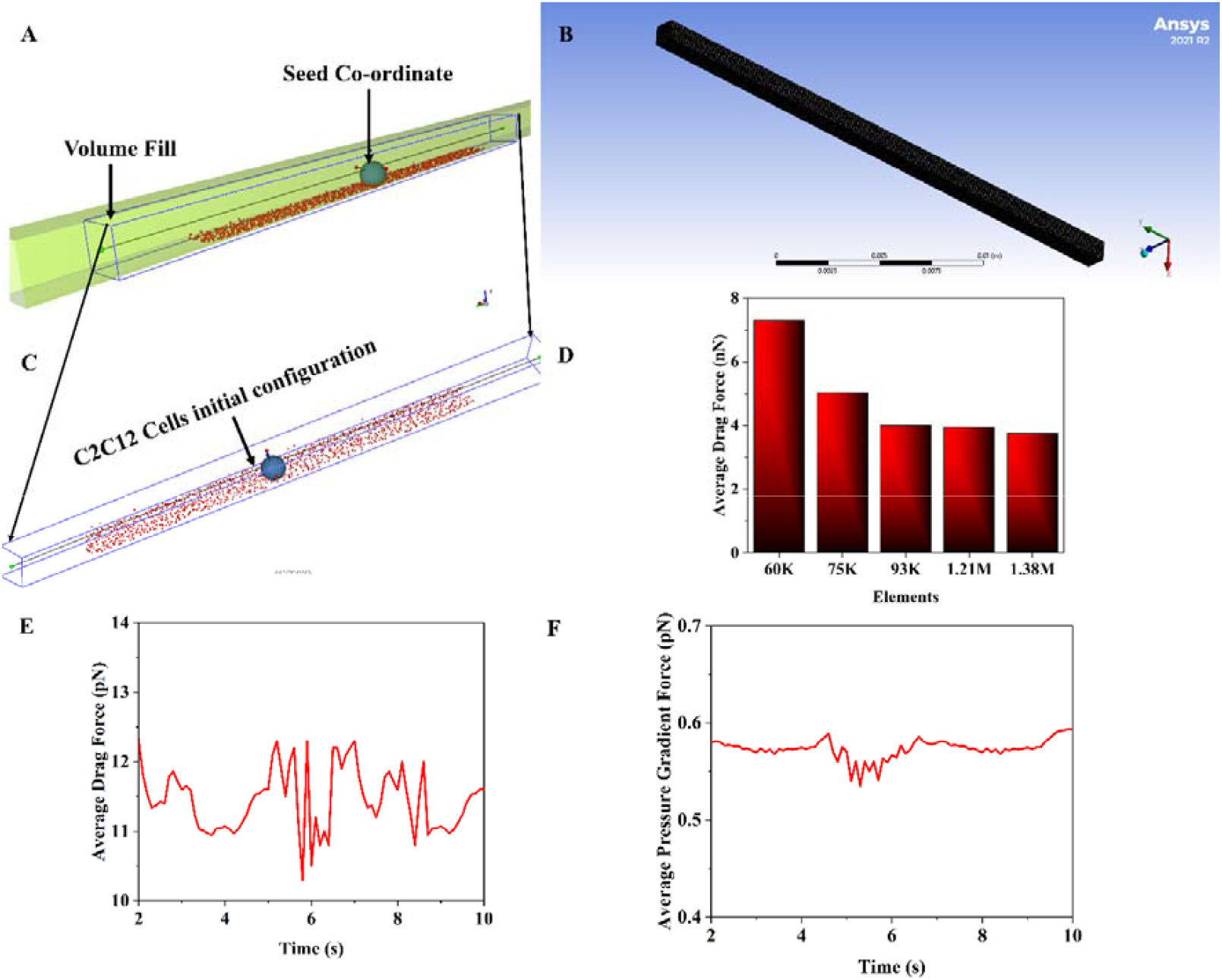
Details of the channel in the microfluidic device used for flow analysis of C2C12 cells. **(a)** CAD geometry of the channel. **(b)** Meshing of the channel. **(c)** Region in the channel where initially C2C12 cells were seeded by volume fill and initial configuration of the cells before flow. The image shows cuboidal box for volume generation and seed co-ordinate which acts as a centre for generation. **(d)** Mesh independent analysis showing variation of average drag force value with number of elements. **(e)** Transient average drag force for flow analysis for 2 hrs of C2C12 cell seeding. **(f)** Transient average pressure gradient force for flow analysis for 2 hrs of C2C12 cell seeding.

In experiments, the images obtained before and after the flow were used to count adhered cells at five different locations along the channel length. An average of these counts was used to compute a dimensionless parameter, termed the Detachment Ratio (DR):

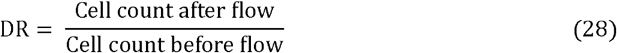

This ratio serves as a quantitative indicator of cell retention post-flow. In simulations, the cell count was intentionally kept lower (917 cells) than in experiments to enable effective tracking and computational efficiency. For the same flow conditions and channel dimensions, simulated DR values closely matched experimental observations, with a deviation of less than 5%, validating the modeling approach.

Once a match was established between simulation and experiment, the simulation data were further used to calculate a dimensionless force parameter, referred to as the Detachment Force Ratio (DFR). DFR quantifies the relative strength of adhesive interactions compared to hydrodynamic forces and is defined as:

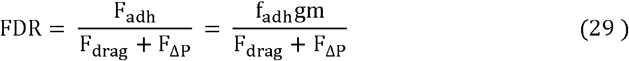

Where, F_drag_ and F_Δp_ are the average fluid induced forces acting on the cells during flow and F_adh_ is the adhesion force. Fig. 12(e) and (f) shows plot for transient average drag force and pressure gradient force for 1 hrs of seeding of C2C12 cells. The average drag force and pressure gradient force is 11.44pN and 0.57pN respectively. In the present study, a constant adhesion force model was employed, wherein the adhesive force is defined as a fixed multiple(f_adh_) of the gravitational force acting on each cell, as detailed in Section 2.5 of the methodology. Whereas the average virtual mass force is significantly less so ignored to calculate the FDR. The single cell mass of C2C12 in the current simulation is Kg and force fraction value is 0.35 in adhesion force model to match DR for 1 hrs which is 58.38% in experiments and near to 60.40% in simulation. Now putting these values in above equation, the FDR value for 2 hrs is calculated as shown below.

Similarly, the calculation was done for all the cases and values are reported in Table 2. Fig. 13 presents the comparison of DR values between simulation and experiment for various seeding times. It is evident from the microscopy images that cell attachment improves with increased incubation time. Furthermore, the post-flow images reveal that the number of retained cells increases with time, indicating stronger bonding due to enhanced interaction between cells and the collagen matrix. Consequently, higher adhesion force values were required in the simulation to accurately capture the experimental trends, especially at longer seeding durations.

**Fig. 13.**
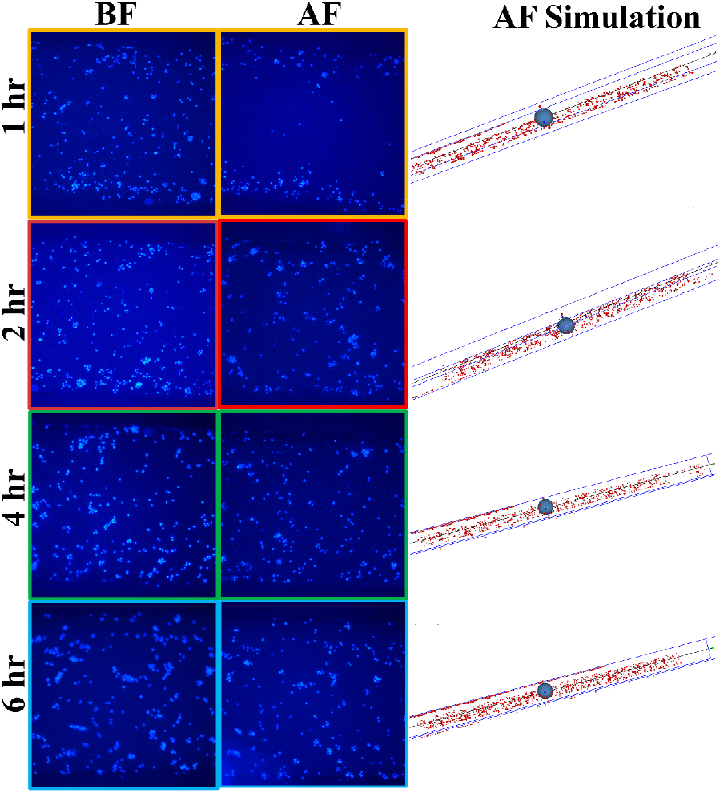
Comparison of the experimental and the simulated data to analyze the adherence of C2C12 cells before and after flow for different seeding time. In the image BF and AF is the abbreviated form of Before Flow and After Flow. The image shows increased cell count after flow with increased seeding time due to strong adhesion between C2C12 cells and collagen matrix.

## 4 Conclusions

This study presents a robust and computationally efficient CFD-DEM model to analyze RBC flow and adherent cell dynamics in a constricted microchannel. To reduce computation time without compromising accuracy, a softening factor as low as 0.03 was implemented, maintaining realistic drag force and cell-free layer (CFL) values. Adhesion between RBCs and the channel wall was incorporated, with a 0.4□µm adhesive interaction range selected based on computational feasibility. RBC–RBC adhesion was excluded due to excessive clustering and CFL distortion.

A coarse-grained model (CGM) with a factor of 1.8 significantly reduced computational load but led to overestimation of drag forces. Rolling resistance (Type C, coefficient 0.8) was introduced to correct this, reducing deviation to within 10% of the original values. The final model was extended to simulate adhesion behavior of C2C12 cells under flow, replicating experiments with varying cell seeding durations. A detachment ratio (DR) was defined based on experimental observations, and the simulation was calibrated accordingly. The detachment force ratio (DFR) reached 0.88 after 6□hours of seeding, indicating strong cell adhesion.

Overall, this optimized model enables accurate simulation of complex microfluidic flows with cell– surface interactions, providing a valuable tool for designing biomedical microdevices and studying cell behavior under physiological conditions.

## Declaration of competing statement

The authors declare that they have no known competing financial interests or personal relationships that could have appeared to influence the work reported in this paper.

